# Sensory-processing traits generate dietary phenotypes that predict microbiome structure: an agent-based modelling framework

**DOI:** 10.64898/2026.09.08.750053

**Authors:** William Ross Hunter

## Abstract

Differences in gut microbiome composition are frequently reported across neurodevelopmental and psychiatric conditions, including autism and depression, and are often interpreted as reflecting underlying neurobiological variation. However, dietary behaviour is both a major determinant of microbiome composition and a downstream consequence of sensory and cognitive processes. To investigate whether microbiome differences can emerge through behavioural pathways alone, I developed a minimal agent-based model linking sensory-processing traits, dietary behaviour, and microbiome structure. Agents were characterised by latent sensory drive, predictability preference, and context sensitivity, which influenced food selection through reinforcement learning and stochastic environmental interactions. Continuous variation in these traits generated a broad spectrum of dietary phenotypes ranging from restrictive to exploratory food-selection strategies. Dietary entropy increased strongly with dietary repertoire size and was associated with reduced microbiome dominance, indicating that more diverse diets produced more even microbial communities. Sensory-processing traits initially appeared to predict microbiome structure directly; however, when dietary entropy was incorporated into the analysis, the direct effects of sensory traits were greatly reduced, while dietary entropy emerged as the dominant predictor of microbiome dominance. Incorporating dietary entropy increased explanatory power from *R²* = 0.67 to *R²* = 0.99. Sensitivity analyses demonstrated that this dietary filtering mechanism was robust across a wide range of assumptions regarding microbiome persistence, reinforcement learning, and choice stochasticity. These findings suggest that dietary filtering provides a plausible mechanism linking sensory-processing characteristics to microbiome structure and may explain at least some microbiome differences reported across gut-brain studies. Integrating sensory, behavioural, dietary, and microbiome data may therefore be essential for improving causal inference within gut-brain axis research.

**Significance Statement:** Differences in gut microbiome composition are widely reported across autism, depression, and other neuropsychiatric conditions. Using a minimal agent-based model, this study shows that variation in sensory-processing characteristics can generate distinct dietary phenotypes that, in turn, produce predictable differences in microbiome structure. Dietary behaviour emerged as the primary mechanism linking sensory phenotype to microbial outcomes, with dietary entropy largely explaining the apparent relationship between sensory-processing traits and microbiome dominance. These findings suggest that some microbiome differences observed in gut-brain research may reflect the downstream consequences of sensory-driven dietary behaviour, highlighting dietary filtering as an important mechanism linking neural and ecological processes.

**Research Highlights:**

- Sensory-processing traits generated stable dietary phenotypes ranging from restrictive to exploratory food-selection strategies within a minimal agent-based model.
- Dietary diversity strongly predicted microbiome structure, with more exploratory diets producing less dominant and more even microbial communities.
- Dietary entropy largely mediated the relationship between sensory phenotype and microbiome dominance, increasing explanatory power from *R²* = 0.67 to *R²* = 0.99.
- The dietary filtering mechanism was robust across variation in microbiome persistence, reinforcement-learning rate, and choice stochasticity.
- Microbiome differences may arise as downstream consequences of sensory-driven dietary behaviour, highlighting the importance of integrating sensory, behavioural, dietary, and microbiome data when interpreting gut-brain relationships.

## Introduction

It has been proposed that the human gut microbiome plays an important role in shaping neurobiology and behaviour (1, 2). Associations between microbiome composition and a range of neurodevelopmental and psychiatric conditions, including autism and depression, are now widely reported (3, 4). These observations are often interpreted as evidence that variation in the microbiome reflects underlying neurobiological differences between individuals (5). However, microbiome associations are frequently inconsistent across studies and rarely map uniquely onto specific diagnostic categories. Similar microbial signatures have been reported across multiple conditions and across the broad phenotypic spectrum of autism itself (6–8). This lack of specificity suggests that microbiome variation may not solely reflect underlying neuropathology, but could instead emerge from behavioural and ecological processes that influence how individuals interact with their environment. Consequently, distinguishing between microbiome patterns that arise from neurobiological mechanisms and those that emerge as indirect consequences of behaviour remains a major challenge for interpreting associations within the gut-brain axis.

This challenge reflects a broader problem in complex biological systems, where similar observable patterns can emerge from the interaction of multiple underlying processes, making it difficult to infer mechanism from outcome alone (9). The problem becomes particularly acute when behaviour acts as an intermediary between biological state and ecological consequence, introducing adaptation, variability and context dependence. Across the gut-brain axis, behaviour represents a pathway through which neurobiological variation can influence microbiome assembly (10). Taking autism as an example, sensory processing differences ranging from hypersensitivity to active sensory-seeking behaviours are widely reported (4). These traits are associated with restricted and selective eating, food neophobia, and heightened sensitivity to food texture and flavour (1, 4). Similar pathways may operate in depression, where changes in appetite, dietary preference and feeding behaviour are commonly observed (2, 3). Given that diet is a major determinant of microbiome composition (10), it is reasonable to expect that sensory and behavioural differences could generate predictable variation in microbial communities. Despite this, behavioural processes are often treated as secondary within microbiome research, with observed compositional differences more frequently attributed to intrinsic biological mechanisms than to the dietary consequences of behaviour (6–8). Consequently, distinguishing microbiome patterns that emerge directly from neurobiology from those arising indirectly through behaviour remains a major challenge for interpreting associations within the gut-brain axis.

I developed a minimal agent-based model that formalises the pathway linking sensory-processing traits, dietary behaviour and microbiome outcomes. Latent sensory characteristics, represented by sensory drive, predictability preference and context sensitivity, influence dietary decisions through reinforcement learning, stochastic choice and environmental modulation. Consequently, dietary phenotypes emerge from behavioural processes rather than being imposed *a priori*. Using this framework, I test whether stable variation in sensory-processing characteristics is sufficient to generate predictable differences in microbiome structure through dietary filtering alone. If so, observed associations between sensory phenotype and microbiome composition need not imply a direct influence of the microbiome on behaviour, but may instead emerge as indirect consequences of long-term dietary patterns. The model is therefore intended as a mechanistic proof-of-concept rather than a quantitatively predictive representation of human dietary or microbial dynamics. Its purpose is not to reproduce specific empirical observations, but to determine whether complex microbiome signatures can emerge from relatively simple behavioural processes operating over extended timescales.

## Results

Dietary behaviour emerged from interactions among latent sensory-processing traits, environmental context and reinforcement learning rather than being prescribed within the model (Fig. 1a). Despite all agents being generated from continuous distributions of sensory drive (S), predictability preference (P) and context sensitivity (C), substantial variation emerged in both dietary repertoire size and dietary entropy (Fig. 1b-c). Consequently, agents organised along a continuum of dietary phenotypes ranging from restrictive food-selection strategies characterised by low dietary entropy to more exploratory phenotypes exhibiting broader and more evenly distributed dietary preferences. Rather than forming discrete behavioural groups, dietary phenotypes occupied a continuous spectrum within the latent sensory space, indicating that relatively small differences in sensory-processing characteristics can generate substantial variation in feeding behaviour.

**Figure 1.**
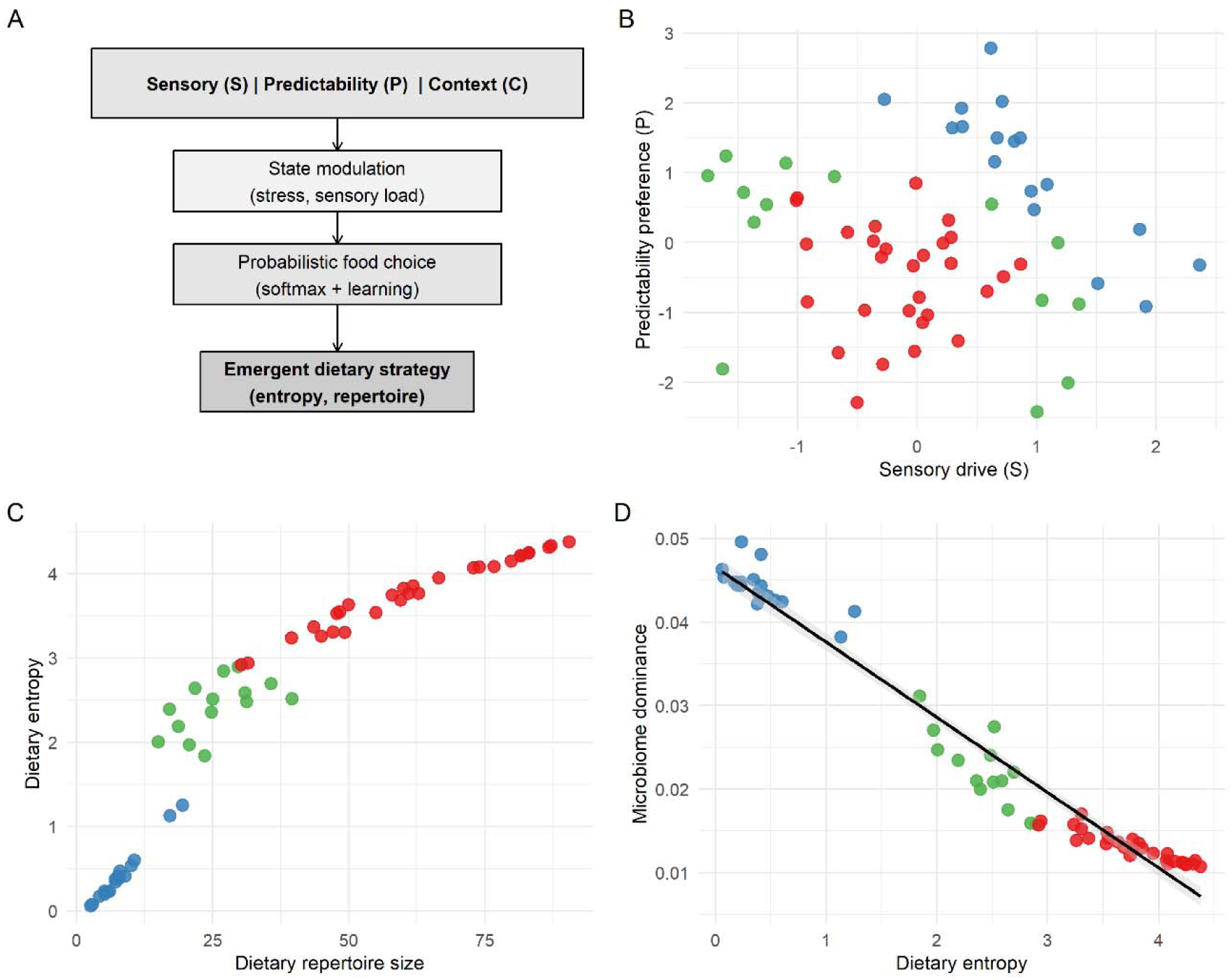
Emergence of dietary phenotypes and microbiome consequences from sensory-processing traits. **a,** Conceptual schematic of the Sensory–Predictability–Context (SPC-Q) model. Latent sensory-processing traits, comprising sensory drive (S), predictability preference (P), and context sensitivity (C), are modulated by environmental state variables (stress and sensory load), influencing food selection through probabilistic decision-making and reinforcement learning. Repeated food choices give rise to emergent dietary phenotypes characterised by dietary entropy and repertoire size. **b,** Latent sensory space defined by sensory drive (S) and predictability preference (P). Points represent individual agents coloured according to emergent dietary phenotype (specialist, opportunistic, or exploratory), illustrating how continuous variation in latent sensory-processing characteristics generates behavioural diversity. **c,** Relationship between dietary repertoire size and dietary entropy across agents. Higher dietary entropy is associated with a broader dietary repertoire, demonstrating the emergence of a continuous gradient of dietary phenotypes ranging from highly restrictive to highly exploratory food-selection strategies. Values represent means across Monte Carlo simulations. **d,** Relationship between dietary entropy and microbiome dominance. Increasing dietary diversity is associated with reduced microbiome dominance, indicating that exploratory dietary phenotypes generate more even microbial communities. Colours correspond to emergent dietary phenotypes shown in Panels B and C. The fitted regression line summarises the population-level relationship across all agents.

Dietary entropy increased strongly with repertoire size across the simulated population (Fig. 1c), indicating that agents consuming a greater number of food types also distributed consumption more evenly across available resources. This relationship emerged directly from the interaction of sensory-processing traits, environmental fluctuations and reinforcement-learning dynamics, demonstrating that structured dietary variation can arise from relatively simple behavioural mechanisms. The resulting dietary phenotypes produced corresponding differences in microbiome organisation (Fig. 1d). Increasing dietary entropy was associated with reduced microbiome dominance, whereas restrictive dietary phenotypes generated communities characterised by a smaller number of dominant taxa. Together, these findings demonstrate that stable differences in sensory-driven dietary behaviour are sufficient to generate predictable ecological differences in microbiome structure within the model framework.

To identify the behavioural mechanisms linking sensory-processing traits to dietary variation, I quantified the effects of sensory drive (S), predictability preference (P) and context sensitivity (C) on dietary entropy (Fig. 2a). All three sensory dimensions were negatively associated with dietary entropy, indicating that stronger sensory constraints reduced dietary diversity. Predictability preference exerted the largest effect (β ≈ −0.52), followed closely by sensory drive (β ≈ −0.49), while context sensitivity exhibited a weaker but still substantial influence (β ≈ −0.23). These findings suggest that a preference for routine and predictability may be at least as important as sensory responsiveness in generating restrictive dietary phenotypes within the model. A similar pattern was initially observed for microbiome structure. When microbiome dominance was modelled directly as a function of sensory-processing traits, all three variables exhibited positive associations with dominance (Fig. 2b). At face value, these results suggest that sensory phenotype directly influences microbial community organisation, with more restrictive sensory profiles associated with communities dominated by a smaller number of taxa. However, this interpretation changed markedly following inclusion of dietary entropy in the model (Fig. 2c). The estimated effects of sensory drive, predictability preference and context sensitivity were reduced to near-zero values, while dietary entropy emerged as the dominant predictor of microbiome dominance (β ≈ −0.86). Consequently, agents exhibiting broader and more evenly distributed dietary patterns consistently developed microbial communities characterised by lower dominance and greater evenness, irrespective of their underlying sensory profile. This shift in explanatory power is consistent with a mediation pathway in which sensory-processing traits influence microbiome structure indirectly through their effects on dietary behaviour. Further support for this interpretation was provided by model performance (Fig. 2d). A model containing only sensory-processing variables explained 67% of the variation in microbiome dominance (R² = 0.67), whereas inclusion of dietary entropy increased explanatory power to 99% (R² = 0.99). Taken together, these findings indicate that sensory-processing traits do not primarily determine microbiome structure directly. Rather, sensory phenotype shapes long-term patterns of food selection, and these dietary phenotypes subsequently influence microbial community organisation through a process of behavioural dietary filtering.

**Figure 2.**
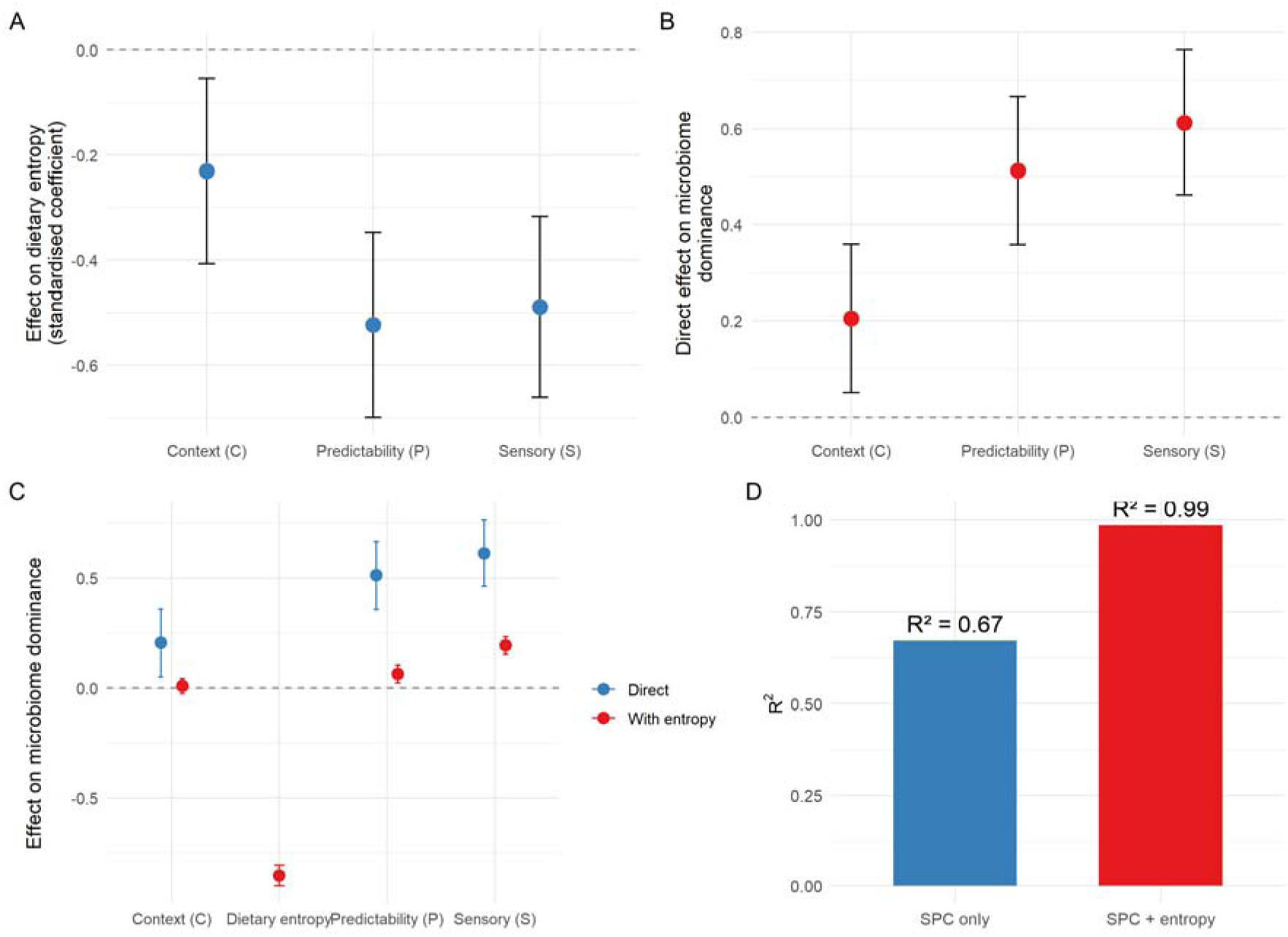
Dietary behaviour mediates the relationship between sensory-processing traits and microbiome structure. **a,** Standardised effects of context sensitivity (C), predictability preference (P), and sensory drive (S) on dietary entropy. Higher predictability preference and sensory drive are associated with reduced dietary diversity, indicating that latent sensory-processing characteristics constrain the breadth and variability of food selection. Points show standardised regression coefficients (β ± 95% confidence intervals). **b,** Direct effects of context sensitivity (C), predictability preference (P), and sensory drive (S) on microbiome dominance. Prior to accounting for dietary behaviour, all three sensory-processing dimensions are positively associated with microbiome dominance, suggesting an apparent relationship between sensory phenotype and microbial community structure. Points show standardised regression coefficients (β ± 95% confidence intervals). **c,** Comparison of direct and mediated models predicting microbiome dominance. Blue points show direct effects of sensory-processing traits, whereas red points show coefficients after dietary entropy is included in the model. Inclusion of dietary entropy substantially attenuates the effects of sensory-processing traits while revealing a strong negative effect of dietary entropy on microbiome dominance, consistent with a mediation pathway linking sensory phenotype to microbiome structure through dietary behaviour. **d,** Variance explained (*R²*) by models predicting microbiome dominance using sensory-processing traits alone (blue) and sensory-processing traits plus dietary entropy (red). Incorporating dietary entropy increases explained variance from *R²* = 0.67 to *R²* = 0.99, demonstrating that dietary behaviour accounts for most of the relationship between sensory-processing traits and microbiome dominance.

To test the robustness of the dietary filtering mechanism, I varied microbiome persistence (λ), reinforcement-learning rate (α), and choice stochasticity (β) across a range of plausible parameter values and quantified the effect of dietary entropy on microbiome dominance (Fig. 3). Across all parameterisations, dietary entropy remained strongly negatively associated with microbiome dominance, indicating that the mediation pathway identified in Figure 2 was not dependent on a specific model configuration. Variation in microbiome persistence produced only minor changes in effect size, suggesting that the relationship was largely insensitive to assumptions regarding microbial memory and turnover. The dietary filtering effect also remained strong across the range of learning rates, although stronger learning modestly increased the magnitude of the entropy-dominance relationship. The largest sensitivity was observed for choice stochasticity. As food selection became increasingly deterministic (higher β), the negative effect of dietary entropy on microbiome dominance strengthened, whereas more stochastic decision-making weakened this relationship. Nevertheless, the effect remained substantial across all parameter combinations considered. These results indicate that microbiome differentiation emerges robustly from dietary filtering and is primarily strengthened by the consistent expression of dietary preferences rather than by specific assumptions regarding microbiome persistence or learning dynamics.

**Figure 3.**
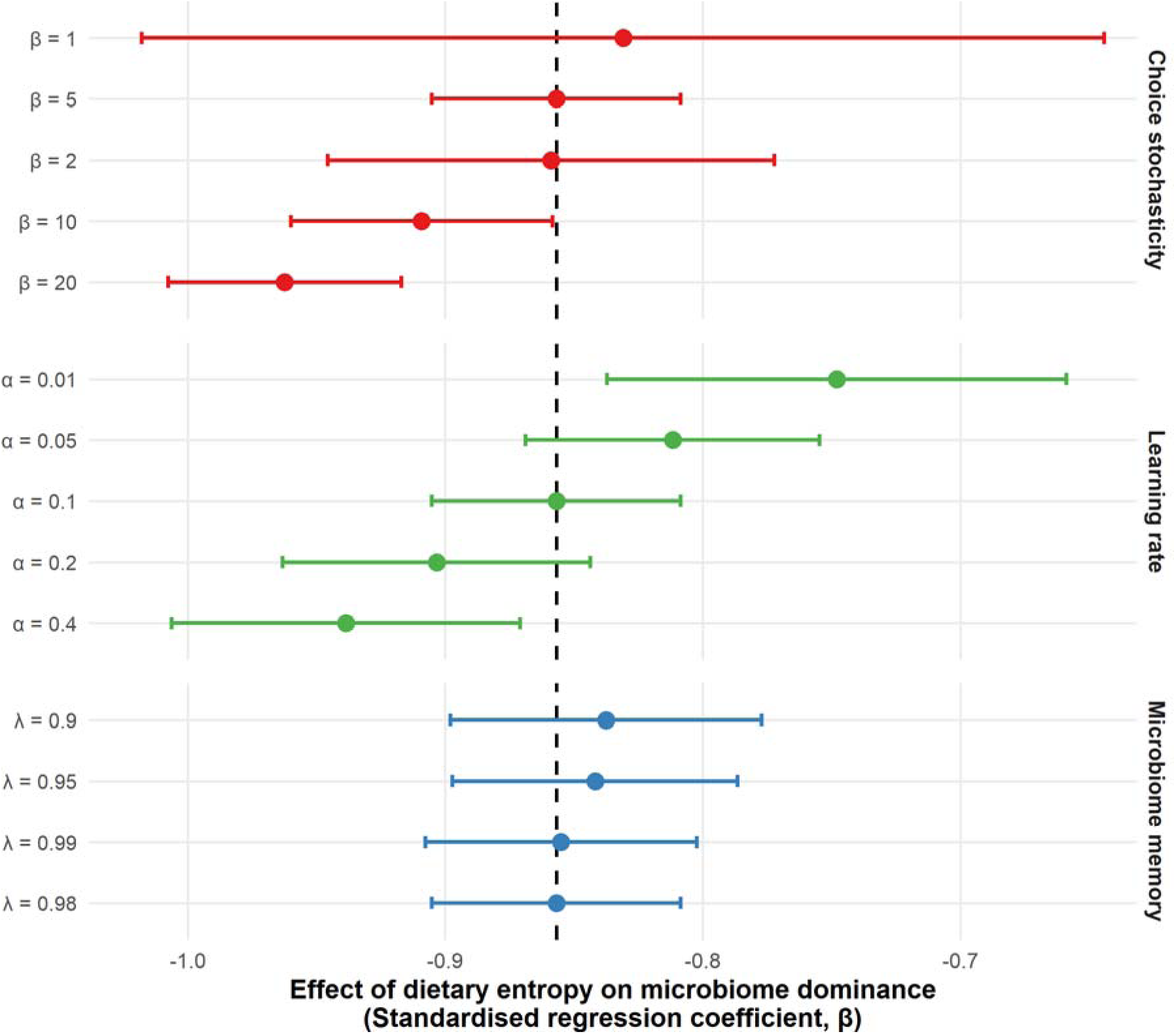
Robustness of dietary mediation across model parameterisations. Points show standardised regression coefficients (β ± 95% confidence intervals) describing the effect of dietary entropy on microbiome dominance under alternative values of microbiome persistence (λ), learning rate (α), and choice stochasticity (β). The dashed line indicates the baseline model estimate (λ = 0.98, α = 0.10, β = 5). Dietary entropy remained a strong negative predictor of microbiome dominance across all parameterisations, indicating a robust mediation effect. Stronger choice determinism increased the effect size, suggesting that microbiome differentiation is enhanced when dietary preferences are expressed consistently through repeated food choices.

## Discussion

Individuals differ substantially in how they perceive and respond to sensory information, how strongly they prefer routine and predictability, and how readily they adapt behaviour to changing environmental conditions. These differences are particularly well documented in autism (1, 4), but are also evident across a broader range of neurodevelopmental and psychiatric conditions characterised by altered sensory processing, appetite regulation and behavioural routine (2, 3). The present study demonstrates that such variation is sufficient to generate distinct dietary phenotypes and, in turn, predictable differences in microbiome organisation. Within the model, sensory drive, predictability preference and context sensitivity shaped long-term patterns of food selection, generating continuous variation in dietary entropy and dietary repertoire. These behavioural differences were strongly associated with microbiome structure, with more diverse dietary phenotypes producing communities characterised by lower dominance and greater evenness. The relationship between sensory-processing traits and microbiome dominance was largely explained by dietary behaviour itself. Although sensory traits initially appeared to predict microbiome structure, these effects were substantially reduced following inclusion of dietary entropy, which emerged as the dominant predictor of microbiome dominance. This pattern is consistent with a mediation pathway in which sensory-processing characteristics influence microbiome organisation indirectly through their effects on food selection. Consequently, the observed microbiome differences did not arise directly from latent sensory traits, but rather from the long-term dietary patterns those traits produced. Together, these findings identify behavioural dietary filtering as a plausible mechanism linking variation in sensory phenotype to predictable differences in microbiome structure (2, 10).

These findings have important implications for the interpretation of gut-brain axis research. Differences in gut microbiome composition have been reported across a range of conditions, including autism, depression, and other neuropsychiatric disorders (3, 4, 6–8). Such patterns are frequently interpreted as reflecting underlying neurobiological differences between individuals. However, diet remains one of the strongest determinants of microbiome composition (10). As such, microbiome structure is likely to be influenced, at least in part, by behavioural and cognitive characteristics that shape long-term dietary exposure. In autism, sensory-processing differences are strongly associated with food selectivity, dietary restriction, and food-related routines (1, 4), while in depression, appetite regulation, motivation, and behavioural routine can similarly influence dietary behaviour (2, 3). The results presented here suggest that at least some differences in microbiome structure may emerge through these behavioural pathways. Consequently, observed microbiome differences should not automatically be interpreted as evidence of microbiome-driven behavioural effects, but may instead reflect the cumulative dietary consequences of underlying sensory and cognitive variation.

A key implication of this study is that behaviour should be viewed as a mechanistic intermediary linking sensory and cognitive processes to ecological outcomes in the gut microbiome. Within the model, dietary choices emerged from interactions among sensory preferences, predictability preferences, environmental context and reinforcement-learning history. Consequently, dietary phenotypes were not direct expressions of latent traits, but emergent properties arising from repeated interactions between individuals and their environment. Because diet acts as a primary resource axis for the microbiome (10), variation in dietary behaviour generated persistent differences in the ecological conditions experienced by microbial communities. In this sense, dietary behaviour acts as a form of environmental filtering, determining which microbial taxa are repeatedly exposed to favourable resource conditions and therefore able to persist and dominate over time.

The gut microbiome represents a dynamically assembled ecological community (11, 12). Community structure emerges through repeated exposure to behavioural resource gradients, analogous to the way environmental conditions structure species assemblages in natural ecosystems (13). The resulting microbial communities, therefore, reflect not only current host characteristics but also the cumulative effects of past behavioural decisions (11, 12). In this sense, microbiome composition exhibits a form of ecological memory, integrating the legacy of historical dietary exposure over extended periods of time (14). This interpretation is consistent with broader evidence linking behaviour, diet and mental health (2), but shifts attention away from static trait-microbiome associations towards the ecological and behavioural processes that generate them. As such, microbial communities may be better understood as ecological records of accumulated behavioural history, shaped by their environment and community assembly, rather than direct markers of latent neurobiological state (10). The ecological interpretation developed here may help explain why similar microbiome signatures are frequently reported across distinct neurodevelopmental and psychiatric conditions. If microbiome composition emerges in part as a response to behavioural effects on dietary exposure, comparable microbial community structures could potentially arise from different combinations of sensory characteristics, behavioural trajectories and dietary environments. Convergence may indicate that similar microbiome compositions do not necessarily indicate common underlying neurobiological mechanisms. Instead reflect comparable ecological histories. This possibility is consistent with the broader principle that similar system-level outcomes can emerge from different underlying pathways (9). However, evaluating that hypothesis would require explicit comparison of alternative behavioural routes to microbiome assembly, which lies beyond the scope of the current model.

This model is relevant for understanding heterogeneity within autism and other neurodevelopmental conditions. For example, Autistic individuals exhibit pronounced variations in sensory sensitivity, sensory seeking, food selectivity, and dietary restriction (4). These characteristics should not be treated as noise around a common diagnosis. Instead, the model suggests that they represent important determinants of dietary phenotype. Figure 1 demonstrates that relatively modest variation in sensory-processing characteristics can produce substantial differences in dietary entropy and repertoire size, while Figure 2 shows that these dietary differences are sufficient to explain variation in microbiome structure. Under this interpretation, microbiome heterogeneity may be an expected consequence of behavioural heterogeneity. This perspective is consistent with the considerable variability and often inconsistent microbial signatures reported across autism cohorts (6–8). Individuals who share a diagnostic label may nevertheless experience markedly different sensory environments and dietary exposures, resulting in divergent microbiome outcomes. More broadly, these findings suggest that variation within conditions may be as important as variation between conditions when interpreting microbiome data (5). Although autism provides the primary motivating example for this study, the mechanism explored here is not autism-specific. Any condition in which sensory processing, behavioural routine, appetite regulation, or food-selection behaviour influence long-term dietary exposure may generate similar microbiome patterns through behavioural dietary filtering. Consequently, the framework may have broader relevance across neurodevelopmental and psychiatric conditions where microbiome differences have been reported.

The sensitivity analysis conducted here demonstrates that the dietary filtering mechanism was robust across a wide range of model parameterisations (Fig. 3). Variation in microbiome persistence (λ) had little influence on the strength of the relationship between dietary entropy and microbiome dominance, indicating that the mediation effect was not dependent on assumptions regarding microbial memory or turnover. Similarly, dietary entropy remained a strong predictor of microbiome dominance across all reinforcement-learning rates (α), although faster learning modestly strengthened the relationship by promoting more stable dietary preferences. The greatest sensitivity was observed for choice stochasticity (β). As food choices became increasingly deterministic, the effect of dietary entropy on microbiome dominance became stronger, whereas more stochastic decision-making weakened this relationship. Overall, dietary entropy remained the dominant predictor of microbiome structure across all parameter combinations examined. Together, these findings suggest that microbiome differentiation emerges primarily through the repeated and consistent expression of dietary preferences rather than through specific assumptions regarding microbiome persistence, learning dynamics, or model parameterisation more generally (15, 16).

The model generates clear empirical predictions regarding dietary filtering as a mechanism linking sensory phenotype to microbiome structure. As such, microbiome variation should covary more strongly with sensory-processing characteristics, food selectivity, and dietary diversity than with diagnostic labels alone. This prediction is consistent with evidence that sensory processing shapes dietary behaviour (4) and with the strong influence of diet on microbiome composition (10). Future studies need to move beyond simple case-control comparisons. Instead, they should collect sensory-processing measures, dietary data, and microbiome sequencing data within the same individuals. In autism research, for example, instruments such as the SPC-Q, Adult/Adolescent Sensory Profile, or Short Sensory Profile could be paired with food-frequency questionnaires, dietary diversity measures, and microbiome profiling. Integrating these behavioural and ecological data would allow direct tests of whether individual differences in sensory processing and dietary behaviour explain variation in microbiome structure more effectively than diagnostic category alone. Such datasets would also provide opportunities to refine model parameters and evaluate the extent to which dietary filtering contributes to observed microbiome differences in real populations.

The model was intentionally developed as a minimal theoretical framework and is not intended as a quantitatively predictive representation of human dietary behaviour or microbiome dynamics. The microbiome module is simplified and does not incorporate many important features of real microbial ecosystems, including interspecific interactions, host physiology, colonisation processes, or feedbacks between the microbiome and behaviour (10, 17). Nevertheless, the simulated microbiome comprised approximately 400 operational taxonomic units (OTUs), comparable to contemporary estimates of individual gut microbial richness in humans (refs). Behaviour was represented solely through dietary choice, whereas real-world dietary patterns are influenced by a wider range of social, economic, cultural, and physiological factors. Consequently, the model should not be interpreted as a realistic simulation of gut microbial ecology. Instead, its purpose was to isolate and examine the behavioural mechanisms capable of generating structured microbiome variation. By reducing the system to a small number of interacting processes, the model demonstrates that dietary filtering alone is sufficient to generate predictable differences in microbiome organisation, providing a theoretical foundation for future empirical and mechanistic studies.

Overall, the model show that stable variation in sensory-processing characteristics can generate distinct dietary phenotypes. In turn, these produce predictable differences in microbiome structure. Within the model, behaviour emerged as the primary mechanism linking latent sensory traits to ecological outcomes, with dietary entropy mediating the relationship between sensory phenotype and microbiome dominance. These findings suggest that dietary filtering provides a plausible explanation for at least some microbiome differences reported across gut-brain studies (2, 4, 10). Rather than viewing behaviour as a nuisance variable or covariate, the results indicate that it should be considered a mechanistically essential component of gut-brain systems. Under this framework, microbiome composition may often reflect the downstream ecological consequences of sensory-driven dietary behaviour rather than direct microbial influences on behaviour itself. Recognising dietary behaviour as a mechanistic intermediary is therefore essential for interpreting microbiome variation, improving causal inference, and developing a more integrated understanding of gut-brain relationships.

## Materials and Methods

### Model Overview

I developed a minimal agent-based model to investigate how dietary behaviour emerges from interactions among latent sensory-processing traits, environmental context, and reinforcement learning (15, 16). Food choice is represented as a stochastic, state-dependent process in which agents repeatedly select from a structured sensory environment (4, 18, 19). Behaviour emerges endogenously through the interaction of stable sensory preferences, contextual modulation, and experience-dependent learning. A microbiome module was incorporated to examine how emergent dietary phenotypes influence downstream ecological outcomes, including microbial community diversity and dominance (10, 17). To account for stochasticity in decision-making and environmental dynamics, simulations were repeated across independent Monte Carlo runs (n = 30) using different random seeds. Monte Carlo replication was used to assess the robustness of emergent behavioural and microbiome outcomes while maintaining a computationally efficient framework appropriate for an illustrative mechanistic model rather than a fully parameterised predictive simulation.

### Agent-based model and decision dynamics

The simulated food environment comprised *N* = 100 discrete food items, each characterised by sensory intensity (*Iᵢ*), novelty (*Nᵢ*), and predictability (*Rᵢ*), with all attributes sampled independently from uniform distributions on the interval [0,1]. A population of *M* = 60 agents was defined, with each agent *a* parameterised by latent sensory drive (*Sₐ* ∼ *N*(0,1)), predictability preference (*Pₐ* ∼ *N*(0,1)), and context sensitivity (*Cₐ* ∼ *U*(0,1)). Sensory drive captures a continuum from avoidance to sensory seeking, predictability preference reflects tolerance for routine versus novelty, and context sensitivity determines the magnitude of environmental modulation of latent states.

Simulations were conducted over *T* = 200 discrete time steps. At each time step *t*, environmental variables, including stress (*σₜ*), sensory load (*Lₜ*), and environmental predictability (*Eₜ*), were sampled independently from uniform distributions on the interval [0,1]. These variables represent short-term contextual fluctuations that modulate, but do not replace, stable sensory-processing characteristics. Agent state was updated according to:

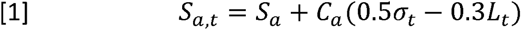

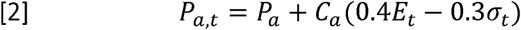

At each time step, agents evaluated available food items using a utility function that integrates current latent state, food attributes, and learned value:

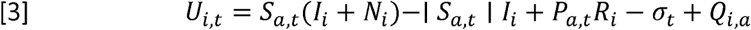

Within this formulation, the term |*Sₐ,t*|*Iᵢ* introduces a penalty for high-intensity foods under both strong sensory-avoidant and sensory-seeking states, capturing nonlinear responses to sensory stimulation during dietary choice. Food choices were generated probabilistically using a softmax decision rule:

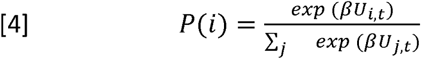

where β = 5 governs the degree of choice stochasticity.

Expected values for each food were updated following each choice according to a standard reinforcement learning rule:

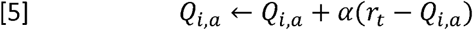

with learning rate α = 0.1 and reward defined as:

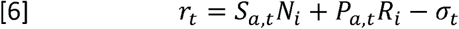

This approach allows dietary preferences to emerge through repeated interactions between stable sensory-processing traits, environmental modulation and accumulated experience.

Dietary behaviour was conceptualised as an emergent dietary phenotype and quantified using two complementary metrics: repertoire size (the number of unique foods selected during a simulation) and dietary entropy:

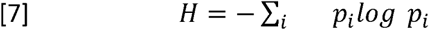

where *pᵢ* denotes the proportion of selections of food *i*. Repertoire size captures dietary breadth, whereas dietary entropy captures both breadth and evenness of food selection. Together these metrics characterise long-term dietary strategy rather than individual food-choice events.

### Microbiome Module

The microbiome module was designed as a simplified representation of how repeated dietary exposures can influence downstream ecological outcomes rather than as a realistic simulation of microbial community dynamics. The simulated microbiome comprised *K* = 400 taxa organised into *n* = 4 functional groups. Each food item was associated with a sparse microbial profile, with responses structured around these functional groups to generate non-uniform mappings between diet and microbiome composition. This abstraction captures the general principle that dietary choices can influence microbial community structure through indirect and highly structured pathways without attempting to represent specific taxa, ecological interactions, or physiological mechanisms.

Following consumption of food *i*, microbiome composition was updated using a leaky-integration process:

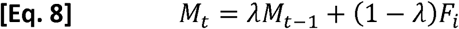

where ***M_t_*** denotes microbiome composition at time *t*, ***F_i_*** represents the normalised microbial profile associated with the selected food item, and ***λ*** determines microbiome persistence (set to 0.98 in the baseline model). Unlike the original implementation, in which microbiome composition was normalised after each update, this formulation allows microbial communities to integrate dietary history over multiple feeding events, producing a cumulative signal that reflects long-term dietary exposure rather than only recent food choices.

At the end of each simulation, microbiome composition was normalised to relative abundance and sampling variation was introduced by drawing multinomial counts (***n*** = **1000**) from the resulting community vector. This procedure generates observed microbial communities analogous to sequencing-derived relative abundance data while preserving the underlying dietary signal.

Microbiome structure was summarised using Shannon entropy, taxonomic richness (number of observed taxa), and dominance (maximum relative abundance), the latter providing an inverse measure of community evenness (20). These metrics were selected to capture complementary aspects of community organisation and to evaluate the ecological consequences of alternative dietary phenotypes.

The microbiome module is intentionally minimal and is not intended to reproduce empirical microbial dynamics or capture key features of real gut ecosystems, including interspecific interactions, host physiology, immigration, extinction, or bidirectional feedbacks between microbial communities and behaviour (10, 17). Instead, it serves as a simplified ecological layer that enables examination of whether stable, sensory-driven dietary phenotypes can generate predictable differences in microbial community structure through dietary filtering alone.

### Dietary phenotype and mediation analysis

To test how sensory-processing traits alter dietary behaviour, agent-level outcomes were averaged across Monte Carlo replicates. Dietary phenotype was characterised using repertoire size and dietary entropy, representing dietary breadth and the diversity of food selection, respectively. Relationships between latent sensory dimensions (sensory drive S, predictability preference P, and context sensitivity C) and dietary phenotype were evaluated using standardised linear models.

To test whether dietary behaviour mediated relationships between sensory phenotype and microbiome structure, microbiome dominance was modelled in two stages. First, microbiome dominance was regressed against latent sensory dimensions (S, P, and C) to estimate direct associations between sensory phenotype and microbial community structure. Second, dietary entropy was added as an additional predictor to evaluate whether dietary behaviour accounted for these associations. Changes in standardised regression coefficients and explained variance (R²) between models were used to assess the extent to which dietary behaviour mediated relationships between sensory-processing traits and microbiome outcomes. Dietary strategies emerged along a continuous gradient of dietary entropy ranging from highly restrictive to highly exploratory food-selection patterns. These behavioural phenotypes were not predefined but emerged from the interaction of latent sensory traits, environmental variation, and reinforcement learning processes operating throughout the simulation.

### Sensitivity analysis

To assess the robustness of the dietary filtering mechanism, sensitivity analyses were conducted across key parameters governing learning, decision-making, and microbiome dynamics. Reinforcement-learning rate (α), choice stochasticity (β), and microbiome persistence (λ) were varied across plausible parameter ranges while all remaining model parameters were held constant. For each parameterisation, simulations were repeated across independent Monte Carlo runs (n = 30) using distinct random seeds. Agent-level behavioural and microbiome outcomes were averaged across Monte Carlo runs prior to analysis to reduce stochastic variability. For each parameterisation, a mediation model relating microbiome dominance to sensory-processing traits (S, P, C) and dietary entropy was fitted. Robustness was quantified using the standardised regression coefficient associated with dietary entropy and its associated 95% confidence interval.

Results were summarised using forest plots comparing effect sizes across parameterisations. This approach follows established agent-based modelling sensitivity-analysis frameworks, in which systematic parameter variation is used to evaluate the stability of emergent mechanisms and model conclusions (15). The primary objective was to determine whether the relationship between dietary entropy and microbiome dominance remained consistent under alternative assumptions regarding behavioural learning, choice dynamics, and microbiome persistence.

### Model implementation and analysis

All simulations and analyses were implemented in R (21). Data manipulation and aggregation were performed using the dplyr and tidyr packages, while visualisation was carried out using ggplot2 (22). Multi-panel figures were assembled using patchwork, model coefficients were extracted using broom, and summary tables were generated using knitr. Monte Carlo simulations were conducted by repeating model runs under different random seeds while retaining the same underlying agents, food environment and food-microbiome mappings. Outputs from individual Monte Carlo runs were combined into a single dataset for analysis. Agent-level summaries were obtained by averaging dietary and microbiome metrics across simulations, allowing stable behavioural phenotypes to be distinguished from stochastic variation in environmental conditions and choice trajectories. Dietary phenotype was characterised using repertoire size, dietary entropy, microbial richness, microbial diversity and microbiome dominance.

Relationships between sensory-processing traits, dietary behaviour and microbiome structure were analysed using standardised linear models. Separate models were fitted to evaluate the effects of sensory drive (S), predictability preference (P), and context sensitivity (C) on dietary entropy, repertoire size and microbiome dominance. Mediation analyses were conducted by comparing models containing only latent sensory-processing variables with models that additionally incorporated dietary entropy. Changes in coefficient magnitude and model explanatory power (R²) were used to assess the extent to which dietary behaviour accounted for relationships between sensory-processing traits and microbiome structure.

Sensitivity analyses were performed to assess the robustness of the dietary filtering mechanism to alternative model parameterisations. Microbiome persistence (λ), reinforcement-learning rate (α), and choice stochasticity (β) were varied systematically across plausible ranges while all other parameters were held constant. For each parameterisation, simulations were repeated using the full Monte Carlo framework (n = 30). Robustness was evaluated using the standardised effect of dietary entropy on microbiome dominance and corresponding 95% confidence intervals.

Visualisations were generated using scatter plots, coefficient plots, and forest plots to illustrate behavioural emergence, mediation pathways, and parameter sensitivity. All figures were exported at publication resolution (300 dpi) using ggsave.

## Code availability

All code used to implement the model and generate the results presented in this study are available at GitHub (https://github.com/Billy-Hunter-AFBI/SPCQ-agent-based-model) and permanently archived on Zenodo (https://doi.org/10.5281/zenodo.20665864).

## Acknowledgments

The author declares no specific acknowledgements.

## Author Contributions

W.R.H. conceived the study, developed the conceptual framework and model, and wrote the manuscript.

## Use of Artificial Intelligence (AI) Tools

The author used Microsoft Copilot to assist with language editing, code annotation, and refinement of reproducible R scripts. All scientific concepts, study design, model development, data analysis, interpretation, and editorial decisions were undertaken by the author, who reviewed and approved all content and retains full responsibility for the accuracy and integrity of the manuscript.

